# Identification and characterization of *ugpE* required for the full virulence of Streptococcus suis

**DOI:** 10.1101/2024.09.25.615082

**Authors:** Qiulei Yang, Na Li, Yanyan Tian, Qiao Liang, Miaomiao Zhao, Hong Chu, Yan Gong, Tong Wu, Shaopeng Wei, He Wang, Guangmou Yan, Fengyang Li, Liancheng Lei

## Abstract

*Streptococcus suis* (*S. suis*) is an emerging zoonotic pathogen that threatens both animal and human health worldwide. UgpE is a protein subunit of the Ugp (<u>u</u>ptake of <u>g</u>lycerol <u>p</u>hosphate) transporter system that is involved in glycerophospholipid synthesis in bacterial membranes. In this study, an *ugpE* deletion mutant was constructed and the effects of *ugpE* deletion on cell morphology, biofilm formation, and virulence were investigated. Deletion of *ugpE* did not affect bacterial growth but impaired cell chain formation and capsular synthesis by downregulating the mRNA levels of the capsular regulon genes *cps-2B*, *cps-2C*, and *cps-2S*. Deletion of *ugpE* also led to decreased tolerance to heat, oxidative, and acid-base stress. Crystal violet staining and scanning electron microscopy demonstrated *ugpE* negatively regulated biofilm formation in liquid culture and the rdar biofilm morphotype on agar plates. Moreover, *ugpE* deletion not only reduced hemolysin activity, survival in whole human blood, and anti-phagocytosis ability against porcine alveolar macrophages (PAM) but also enhanced bacterial adhesion and invasion of human cerebral microvascular endothelial cells (hCMEC/D3) by upregulating the expression of multiple genes associated with cell adhesion. In a mouse infection model, *ugpE* deletion significantly attenuated virulence and lowered the number of viable bacteria in the blood and major organs, as well as distribution of macrophages. In conclusion, this study identified UgpE as a novel virulence factor that plays a pivotal role in the regulation of virulence and biofilm formation of *S. suis*.

## Introduction

*Streptococcus suis* (*S. suis*) is an important zoonotic pathogen that causes meningitis, septicemia, endocarditis, arthritis, and pneumonia in pigs, leading to huge economic losses to the pig industry. *S. suis* also infects humans through direct contact with pathogen-harboring droplets or wounds, causing diseases such as meningitis and toxic shock syndrome (STSS) that threaten human health. *S. suis* is classified into 29 serotypes (1-34, 1/2) according to its capsule antigen, among which *S. suis* serotype 2 (SS2) is the most prevalent and pathogenic strain in many countries, including China, Canada, and Brazil [1, 2]. In 1998 and 2005, outbreaks of human SS2 infection resulted in dozens of deaths in China [3]. In addition, SS2 has become the main pathogen causing meningitis in adults in southeast Asian countries and regions such as Vietnam, Thailand and Hong Kong [4, 5, 6].

The pathogenicity of *S. suis* depends on its virulence factor. To date, a variety of *S. suis* virulence factors have been identified, including capsular polysaccharide (CPS) [7], extracellular factor (EF) [8], muramidase-released protein (MRP) [9], suilysin (SLY) [10], glutamate dehydrogenase (GDH) [11], and fibronectin-binding protein (FbpS) [12, 13]. However, some experimental results indicate that the absence of one or more of these virulence factors does not significantly affect the virulence of certain *S. suis* strains, suggesting that some of these virulence factors may act synergistically in pathogenicity or that new virulence factors may be discovered. Therefore, there are still many unsolved aspects of the pathogenesis of *S. suis* that need to be further investigated to better prevent and control its associated diseases with *S. suis*.

ATP-binding cassette (ABC) transporters constitute a ubiquitous superfamily of integral membrane proteins that are responsible for the ATP-powered translocation of many substrates across membranes. Thus, they play vital roles in the regulation of a variety of physiological processes, including nutrient uptake, drug resistance, lipid transport, and the secretion of non-classical signaling molecules [14]. In prokaryotes, ABC transporters can take up nutrients, efflux toxins, and drugs, thereby affecting bacterial growth and enabling bacteria to resist various antibiotics. In addition, they play an important role in the interactions between pathogens and hosts. They are regarded as virulence factors in uropathogenic *Escherichia coli* because of their critical role in pathogenesis [15]. In *Streptococcus pneumoniae*, the ABC transporter BacA is required for the maintenance of chronic infections in mice [16]. In *S. suis*, proteome analysis has identified genes encoding diverse ABC transporters that may play roles in colonization and infection processes [17, 18, 19, 20]. Moreover, the ABC transporter component ATPase MsmK regulates the pathogenicity of *S. suis* by utilizing carbohydrates [21]. ABC transporters are also involved in the regulation of biofilm formation in various streptococci [22].

The uptake of glycerol phosphate (Ugp) system belongs to the ABC transporter superfamily and is involved in glycerophospholipid synthesis by uptake of *sn*-glycerol-3-phosphate (G3P) and certain glycerophosphodiesters from the periplasm to the cytosol in *E. coli* [23]. The Ugp transporter consists of four subunits known as the UgpABCE system, of which UgpA and UgpE constitute the transmembrane domains, UgpC forms a homodimer of the ATP-hydrolyzing subunit, and UgpB is a periplasmic substrate-binding protein [24]. In addition to glycerophospholipid synthesis, Ugp transporters are also involved in the regulation of other cellular functions by the uptake of G3P and glycerophosphodiesters. Glycerophosphodiesters serve as an osmoprotectant in eukaryotic cells [25, 26, 27]. Moreover, glycerophospholipids are required for the normal growth and virulence of *Mycoplasma pneumoniae* [28, 29]. However, the role of Ugp transporters in the regulation of virulence in other bacterial species remains unknown.

In this study, we assessed the role of *ugpE* (gene ID: YP_003028379.1), a homologous gene that encodes the Ugp transporter subunit UgpE (protein ID: ARL70172.1), in the regulation of various cellular phenotypes, including growth, stress tolerance, biofilm formation, and virulence of the zoonotic pathogen SS2. Our study contributes to elucidating the molecular mechanisms underlying the pathogenesis of *S. suis* and to the development of novel therapies and vaccines against diseases caused by *S. suis*.

## Materials and methods

### Bacterial strains and culture conditions

All strains used in this study are listed in Table S1. SS2 strain SC19 was kindly provided by Prof. Anding Zhang (Huazhong Agricultural University, Wuhan, China). Bacterial cultures were performed under different conditions according to the experimental requirements. SS2 and its derivatives were cultivated in Todd-Hewitt broth (THB; Hopebio, HB0311-3) or plated on THB agar plates supplemented with kanamycin (100□µg/mL; Beyotime, ST101) and spectinomycin (100□µg/mL; Macklin, S6106), if relevant. *E. coli* strain DH5α was cultured in Luria-Bertani (LB; Becton Dickinson, DF0446-17-3) liquid medium or on LB agar plates containing 100□µg/mL spectinomycin if relevant.

### Construction of mutant and complemented strains

The *ugpE* chromosomal deletion mutant was constructed by homologous recombination as previously reported [30]. Briefly, the upstream and downstream homologous regions of *ugpE* were amplified using the A1/A2 and A3/A4 primer pairs, resulting in two fragments that were fused and ligated into the pSET4s shuttle vector [31]. The recombinant plasmid was electroporated into SC19 cells, and the bacteria were selected on THB agar plates supplemented with 100 μg/mL spectinomycin at 37°C. The mutant strains (Δ*ugpE*) were confirmed by spectinomycin susceptibility tests and PCR, using the primer pair A5/A6. Gene complementation was conducted by cloning *ugpE* into the pSET2 shuttle vector [31] using primer Y1/Y2, which contains a putative promoter region. The inserted DNA sequences were confirmed using PCR and DNA sequencing. The recombinant plasmid was electroporated into the mutant strain to produce the complementary strain CΔ*ugpE*. The primers and plasmids used are listed in Tables S1 and S2, respectively.

### Growth curves

Briefly, *S. suis* (WT, Δ*ugpE*, and CΔ*ugpE*) cells were cultured overnight, diluted 1:100, and grown to the exponential phase in THB (with 10% FBS). The bacteria were diluted in THB again at 1:100 dilution and incubated at 37°C under vigorous shaking at 180 rpm. The OD_600_ value was measured every hour for 10 h. Growth curves were plotted using GraphPad Prism (version 9.0; San Diego, CA, USA).

### Observation of colony morphology

The bacterial chain length and capsule were observed using light and transmission electron microscopy (TEM), respectively. Briefly, a suspension of bacterial cells (WT, Δ*ugpE*, CΔ*ugpE*) grown overnight was diluted in THB (with 10% FBS) medium and were diluted in THB again at 1:100 dilution and incubated at 37°C under vigorous shaking at 180 rpm. For light microscopy, the cells were harvested at the mid-log growth phase for Gram staining. The bacterial chain length was observed under a light microscope (Nikon) and determined by counting the cell numbers of 200 chains per cell. For TEM, cells in the mid-log growth phase were collected and fixed in 2.5% glutaraldehyde overnight. After dehydration with propylene oxide and embedding in epoxy resin, the samples were sectioned and stained with 1% uranyl acetate and alkaline lead citrate and observed under a Hitachi HT 7700 (Hitachi, Tokyo, Japan) electron microscope at 80□kV. The bacterial cell width, cell wall, and capsule thickness were determined using ImageJ software (version 1.8.0) by randomly selecting 20 cells.

### Bacterial stress tests

Bacterial stress tests were performed as previously described with slight modifications [32]. Briefly, overnight cultures of *S. suis* cells (WT, Δ*ugpE*, CΔ*ugpE*) were diluted in THB medium at 1:100 dilution and grown to mid-log stage at 37°C. For the hydrogen peroxide (H_2_O_2_) test, cells were diluted again to OD_600_ = 0.1 and grown to the mid-log stage. H_2_O_2_ (30 mM) and PBS (control) were added to the cells and incubated at 37°C for 20 min. For the heat stress analysis, cells were also diluted to OD_600_ = 0.1 and incubated at 37°C and 40°C for 12 h. For the acid tolerance test, cells were harvested after centrifugation at 4°C, washed three times with PBS (pH 7.4), and cultured with fresh THB medium at different pH values (2.0, 4.0, 6.0, 6.0, 7.0, 8.0, 10.0, and 12.0) for 45 min at 37°C. For all assays, cells were plated on THA plates after serial dilution in PBS and bacterial numbers were counted after overnight incubation. Each sample was set up in triplicate and repeated independently three times.

### Bacterial adhesion and invasion assays

Adhesion and invasion assays were performed in the human cerebral microvascular endothelial cell line, hCMEC/D3, as described previously [33]. hCMEC/D3 cell line was kindly provided by Prof. Yan Chen (College of Life Sciences, Jilin University). Briefly, hCMEC/D3 cells were cultured to approximately 2.5 × 10^5^ cells/well in a 24-well plate, followed by infection with different groups of bacteria at a multiplicity of infection (MOI) of 100:1. After incubation at 37°C and 5% CO_2_ for 1 h, the cells were washed four times with PBS to remove non-adherent bacteria. For the adhesion assay, cells were lysed with 0.1% saponin. Bacterial adherence was calculated as follows: (recovered CFU/initial inoculum CFU) × 100%. For the invasion assay, cells were incubated with fresh DMEM supplemented with 5 μg/mL penicillin G (Sangon Biotech; B540729-0010) and 100 μg/mL gentamicin (Sangon Biotech; E607063-0100) for 2 h to kill non-adherent bacteria and then lysed with 0.1% saponin (Merck; 47036). Lysates were plated on THA plates after serial dilution in PBS and bacterial numbers were counted after overnight incubation. Bacterial invasion was calculated as follows: (recovered CFU/adhered CFU) × 100%. Each sample was set up in triplicate and repeated independently three times.

### Phagocytosis

The phagocytosis assay was conducted in porcine alveolar macrophages (PAM) as described previously, with slight modifications [34]. Immortalized PAM (ATCC, CRL-2845) was provided by the Harbin Veterinary Research Institute of the Chinese Academy of Agricultural Sciences. PAM cells were cultured to approximately 2.5 × 10^5^ cells/well in a 24-well plate, followed by infection with different groups of bacteria at a MOI of 40:1. After incubation at 37°C and 5% CO_2_ for 1 h, the cells were washed three times with PBS and incubated with fresh DMEM supplemented with 5 μg/mL penicillin G and 100 μg/mL gentamicin for 2 h to kill non-adherent bacteria. The cells were then washed three times with PBS and lysed with trypsin (ThermoFisher; R001100). Lysed cells were serially diluted and plated on THA plates for counting. Each sample was set up in triplicate and repeated independently three times.

### Whole blood survival

Each group of bacteria (WT, Δ*ugpE*, CΔ*ugpE*) was cultured in THB supplemented with 1% glucose until mid-log growth phase. After centrifugation, the bacterial body was collected and resuspended to an OD_600_ of 0.2 in PBS. Then, 1 mL of germ-free human whole blood containing EDTA-Na_2_ anticoagulant was added to 100 μL of the bacterial samples and incubated at 37°C for 2h. The mixture was serially diluted and plated on THA plates for cell counting. Survival was calculated as follows: (recovered CFU/CFU in the original inoculum) ×100%. The experiment was repeated three times, and three samples were collected in triplicate each time.

### Biofilm formation

Biofilm formation (adherence to the well wall) was assessed as described previously, with slight modifications [35]. Briefly, each group of bacteria (WT, Δ*ugpE*, CΔ*ugpE*) was incubated in 96-well plates with 200 μL THB medium per well at 37°C for 24h, 48h, and 72h. After removing the unattached bacteria, adherent cells were stained with 0.2% crystal violet, washed, and dissolved in a 95% alcohol solution. Biofilms were measured as the absorbance of dissolved crystal violet at OD_595_. Rdar (red, <u>d</u>ry, <u>a</u>nd rough) morphotype assessment was performed as described previously [36]. Briefly, 5□µL of an overnight culture of OD_600_□=□5 suspended in water was spotted onto THA plates containing the dye Congo red (40□µg/mL) and Coomassie Brilliant Blue G-250 (20□µg/mL) or calcofluor white (Fluorescence Brightener 28, 50□µg/ml). After incubation at 37°C for 24h, 48h, and 72h, pictures were taken to analyze the development of the colony morphology structure and dye binding. The surface morphology of the colonies was plotted and analyzed using the ImageJ software (version 1.8.0).

### Scanning electron microscopy

Scanning electron microscopy (SEM) was performed to visualize the macrocolony biofilms. An overnight culture of OD_600_□=□5 suspended in water was prepared and spotted on THA plates pre-embedded with a rectangular glass slide. After 48h of culture at 37°C, the macrocolonies on the glass slide were fixed in fixative solution (0.5% glutaraldehyde and 2.5% paraformaldehyde in 10□mM HEPES, pH 7.0), dehydrated with acetone, sputter-coated with gold/palladium, and observed using a Zeiss Merlin field emission SEM at an acceleration voltage of 10□kV with an Everhart-Thornley secondary emission (SE) detector.

### RNA isolation and qRT-PCR

Each group of bacterial cells (WT, Δ*ugpE*, CΔ*ugpE*) was grown in THB medium until the mid-log growth phase. Total RNA was extracted using TRIzol reagent (Invitrogen, USA), according to the manufacturer’s instructions. RNA was treated with DNase (Invitrogen, USA), and RNA quality was assessed by gel electrophoresis and PCR. The RNA concentration was measured using a NanoDrop 2000 system (Thermo Scientific). cDNA was synthesized using 1 μg of RNA from each sample using the Prime Scrip RT Master Mix Kit (TaKaRa, Dalian, China). Quantitative PCR (qPCR) was performed using TB Green PCR Master Mix (TaKaRa, Dalian, China) on a Quantagene q225 qPCR System (Kubo, Beijing, China). The 16S rRNA gene was used as an internal reference gene for normalization. Data were analyzed using the 2^−ΔΔCT^ method. All experiments were performed at least three times. The primers used are listed in Table S2.

### Animal experiments

All animal experiments were approved by the Institutional Animal Care and Use Committee of Jilin University and were conducted in accordance with the Chinese Laboratory Animal Administration Act 1988 and the Helsinki Declaration of 1975. To determine bacterial virulence in mice, forty 6-weeks old female BALB/c mice (body weight 18 ± 2 g, purchased from Changsheng Biotechnology Co., Ltd., China) were divided into four groups (10 mice per group) randomly and intraperitoneally infected with 100 μL (1×10^9^ CFU, LD_50_=5×10^8^ CFU) of SC19, Δ*ugpE*, CΔ*ugpE* or PBS (as control). All mice were housed in standard plastic mouse cages and kept under constant room temperature (23 ± 3°C), humidity (55 ± 5 %), and observed for one week. Clinical scores, body weight changes, and animal deaths were recorded every 12 h.

For histopathological analysis, mice were intraperitoneally infected with 100 μL (5×10^8^ CFU) of each strain or PBS (as a control). After 3 days, the mice were euthanized by inhalation of CO_2_ and organs including the brain, lung, liver, spleen, and kidney were extracted and frozen at −80°C or used immediately for histopathological analysis and flow cytometry. Histopathological analysis was performed as described previously [37]. Briefly, the tissues were fixed in 10% formaldehyde solution, embedded in paraffin, sliced, and stained with hematoxylin and eosin (HE). The stained sections were visualized, and pathological changes in the tissues were analyzed under an optical microscope (Olympus, Tokyo, Japan). The histological score was assessed as previously described [38]. Additionally, to evaluate the distribution of bacteria in different organs after infection, equal weights of brain, lung, liver, spleen, kidney, and blood samples were homogenized, diluted, and plated on THA plates for cell counting.

### Flow cytometry

Macrophages in the lung, liver, spleen, heart, and kidney were assessed using flow cytometry, as previously reported [39]. Briefly, the organs were shred and digested in HBSS containing gelatinase A (100 U/mL) and DNase (20 µg/mL). After filtration and rinsing with PBS, the samples were centrifuged and the pellets were treated with ice-cold RBC lysis buffer for 5 min. The isolated cells were washed with ice-cold PBS again and centrifuged to collect the cell pellets for antibody labeling in a fluorescent washing buffer (Thermo Fisher). The cell suspension was incubated with BV421 anti-mouse CD11b, APC anti-mouse F4_80, PE anti-mouse CD206, PE-Cy5.5 anti-mouse CD45, and PE-Cy7 CD86 anti-mouse antibodies in the dark for 30 min. After centrifugation, washing with PBS, and resuspension in FWB, the same number of cells in each group was processed with a flow cytometer (CytoFLEX) and analyzed with FlowJo software (Version 10.4).

### Statistical analysis

The experimental data were statistically analyzed using GraphPad Prism (version 9.0, San Diego, CA, USA). Data are presented as mean□±□SD from three independent replicates. Differences between the mean values of normally distributed data were assessed using one-way ANOVA (Dunnett’s test). *P*-value < 0.05, as indicated by “*”. *P*-value < 0.01 indicated that the difference was extremely significant, as indicated by “**.”

## Results

### ugpE deletion inhibited chain formation and capsular synthesis of SS2

To investigate the role of *ugpE* in SS2, an isogenic *ugpE* deletion mutant strain (Δ*ugpE*) and a corresponding revertant strain (CΔ*ugpE*) were constructed by homologous recombination (Fig. S1). We first examined the effect of *ugpE* on the growth kinetics. Although the Δ*ugpE* mutant appeared to grow more slowly than the SC19 wild-type, as observed by the growth curve, the differences between each group were not statistically significant (Fig. 1A). Interestingly, observation of cell morphology by Gram staining and light microscopy revealed that the Δ*ugpE* mutant displayed abnormal cell chain formation compared to the wild-type (Fig. 1B and C). After 72h of culture in THB medium, the majority of the bacterial chains of the SC19 wildtype consisted of 1–4 cell (s) per chain (approximately 80%), while the rest consisted of more than 5 cells per chain, and some even displayed very long chains with more than 10 cells, which was not observed in the Δ*ugpE* mutant (Fig. 1C; Fig. S2). On the other hand, most of the Δ*ugpE* mutants consisted of only 1–3 cell (s) (approximately 95%), and bacteria consisting of more than four cells only accounted for a minor proportion. The majority of the revertant strains also consisted of 1–4 cell (s) (approximately 70%), but no cells with long chains (>10 cells per chain) were found (Fig. 1C; Fig. S2). Moreover, deletion of *ugpE* also led to a significantly decreased cell width compared to the SC19 wildtype, as confirmed by transmission electron microscopy (TEM) (Fig. 1D and E). Overall, these results suggest a role for *ugpE* in regulating SS2 chain formation.

**Fig. 1.**
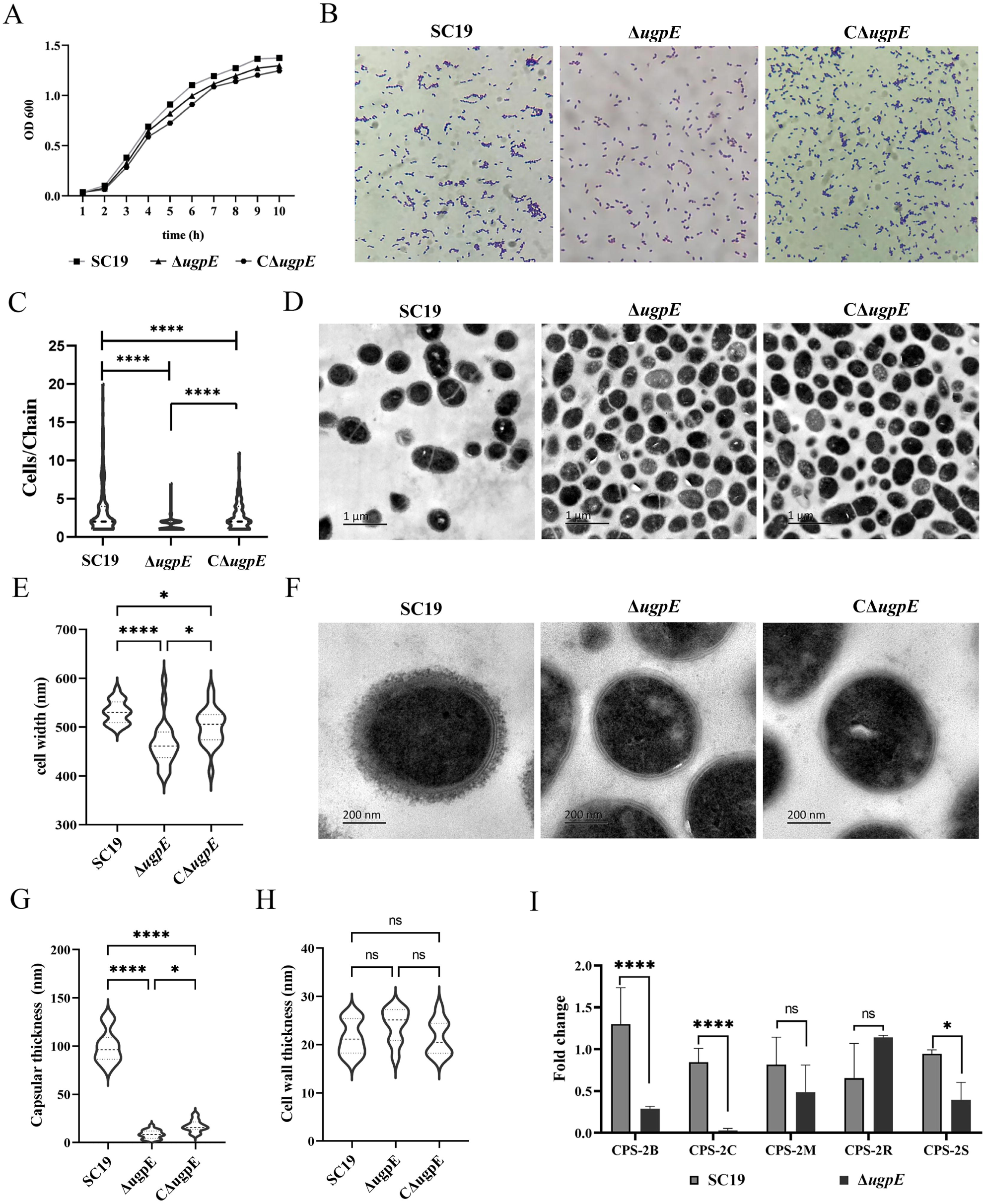
*ugpE* deletion inhibited chain formation and capsular synthesis of SS2. (A) Growth kinetics of SC19 wildtype, Δ*UGPE* and CΔ*UGPE* at 37°C. (B) Observation of cell morphology by Gram staining and light microscopy (1000×). (C) Analysis of cell number per chain in each group of cells. The quantification is based on results from at least three independent experiments with the assessment of 200 cells from each group. (D) Observation of cell morphology by transmission electron microscopy (TEM). Size bar, 1 μm. (E) Analysis of cell width in each group of cells. The quantification is based on results from at least three independent experiments with the assessment of 20 cells from each group. (F) Observation of capsule structure by TEM. Size bar, 200 nm. (G) Analysis of capsular thickness in each group of cells. The quantification is based on results from at least three independent experiments with the assessment of 20 cells from each group. (H) Analysis of mRNA levels of cps regulon genes *cps-2B*, *cps-2C*, *cps-2S*, and *cps-2M* by qRT-PCR. The data are presented as mean ± SD. Data were analyzed using one-way ANOVA (Dunnett’s test) (C, E, G) and paired *t* tests (H), respectively. \*\*\**P* < 0.001; \*\**P* < 0.01; \**P* < 0.05. ns, not significant.

Of note, TEM also showed that the SC19 wild-type had a thick capsule structure, while the capsule structure of Δ*ugpE* was hardly observed (Fig. 1F). In line with this, the capsular thickness of the Δ*ugpE* strain was significantly thinner than that of the SC19 and revertant strains (Fig. 1G). In contrast, cell wall thickness showed no obvious differences among the groups (Fig. 1H). Knowing that the bacterial capsule structure was abolished, we assessed the level at which *ugpE* affected the capsular regulon genes. The *cps* gene cluster of SS2 comprises 25 genes [40], and we chose several of them for qRT-PCR analysis. Compared to the SC19 wildtype, the mRNA levels of *cps-2B*, *cps-2C*, and *cps-2S* were downregulated, while the mRNA levels of *cps-2M* and *cps-2R* were not affected (Fig. 1I). Overall, these results indicate that *ugpE* is required for capsule synthesis of SS2.

### ugpE deletion led to decreased tolerance to environmental stresses

In a series of processes, such as survival and pathogenesis, pathogens have to face various stresses from constantly changing environments, such as acid-base tolerance and hydrogen peroxide. The anti-stress ability directly determines bacterial survival; thus, we determined the bacterial anti-stress ability by exposing it to various stress challenges, including heat stress, hydrogen peroxide killing, and acid-base tolerance. As shown in Fig. 2A, the number of recovered viable bacteria in the Δ*ugpE* mutant was significantly lower than in the SC19 wildtype at both 37°C and 40°C (Fig. 2A). Similarly, the Δ*ugpE* mutant was more sensitive to hydrogen peroxide (30 mM) than the SC19 wildtype and revertant strain (Fig. 2B), indicating that the antioxidant capacity of SS2 was downregulated after *ugpE* deletion. Moreover, the acid-base tolerance test showed that the survival ability of the Δ*ugpE* mutant was significantly lower than that of the SC19 wild-type and revertant strains at all indicated pH points, except at pH8, where no differences were observed between the groups (Fig. 2C). Overall, these results suggest that *ugpE* contributes to bacterial resistance to high-temperature adaptability, hydrogen peroxide, and acid-base tolerance.

**Fig. 2.**
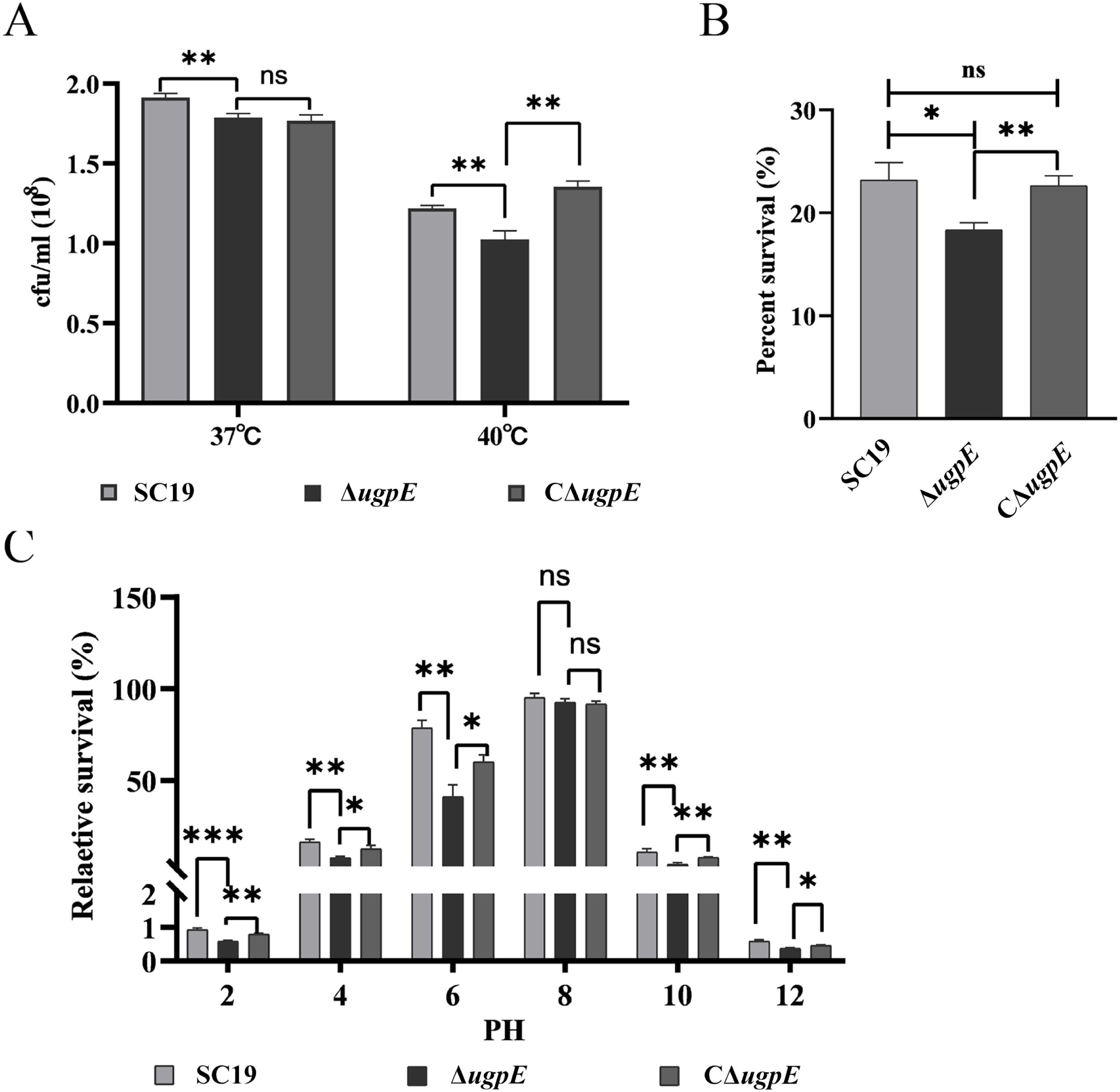
*ugpE* deletion led to decreased tolerance to stresses. (A) Heat stress assay of SC19 wildtype, Δ*ugpE* and CΔ*ugpE* at 37°C and 40°C. (B) Oxidative stress assay of each group of cells. Cells were treated with hydrogen peroxide (30 mM). (C) Acid-base stress assay of each group of cells. The data are presented as mean ± SD. Data were analyzed using one-way ANOVA (Dunnett’s test). \*\*\**P* < 0.001; \*\**P* < 0.01; \**P* < 0.05. ns, not significant.

### ugpE deletion upregulated SS2 biofilm formation

Biofilm formation represents a protective mode of growth that enables pathogens to survive in hostile environments [41]. Since deletion of *ugpE* led to decreased anti-stress ability, we wondered if biofilm formation by SC19 was also affected by *ugpE.* To this end, each group of strains was cultured on 96-well polystyrene plates for 24h, 48h, and 72h, and then stained with crystal violet to visualize biofilm formation. We observed that the biofilms were mainly formed as a ring at the air-liquid interface (Fig. 3A). Interestingly, the biofilms of the Δ*ugpE* mutant displayed a significantly higher level of adherence to the abiotic polystyrene wall of the well surface after 24h, 48h, and 72h of incubation in standing culture at 37°C compared to the SC19 wild-type and revertant strains (Fig. 3A and B). We also detected the effect of *ugpE* on rdar (red, <u>d</u>ry, <u>a</u>nd rough) macrocolony morphotypes using a Congo red agar plate assay.

**Fig. 3.**
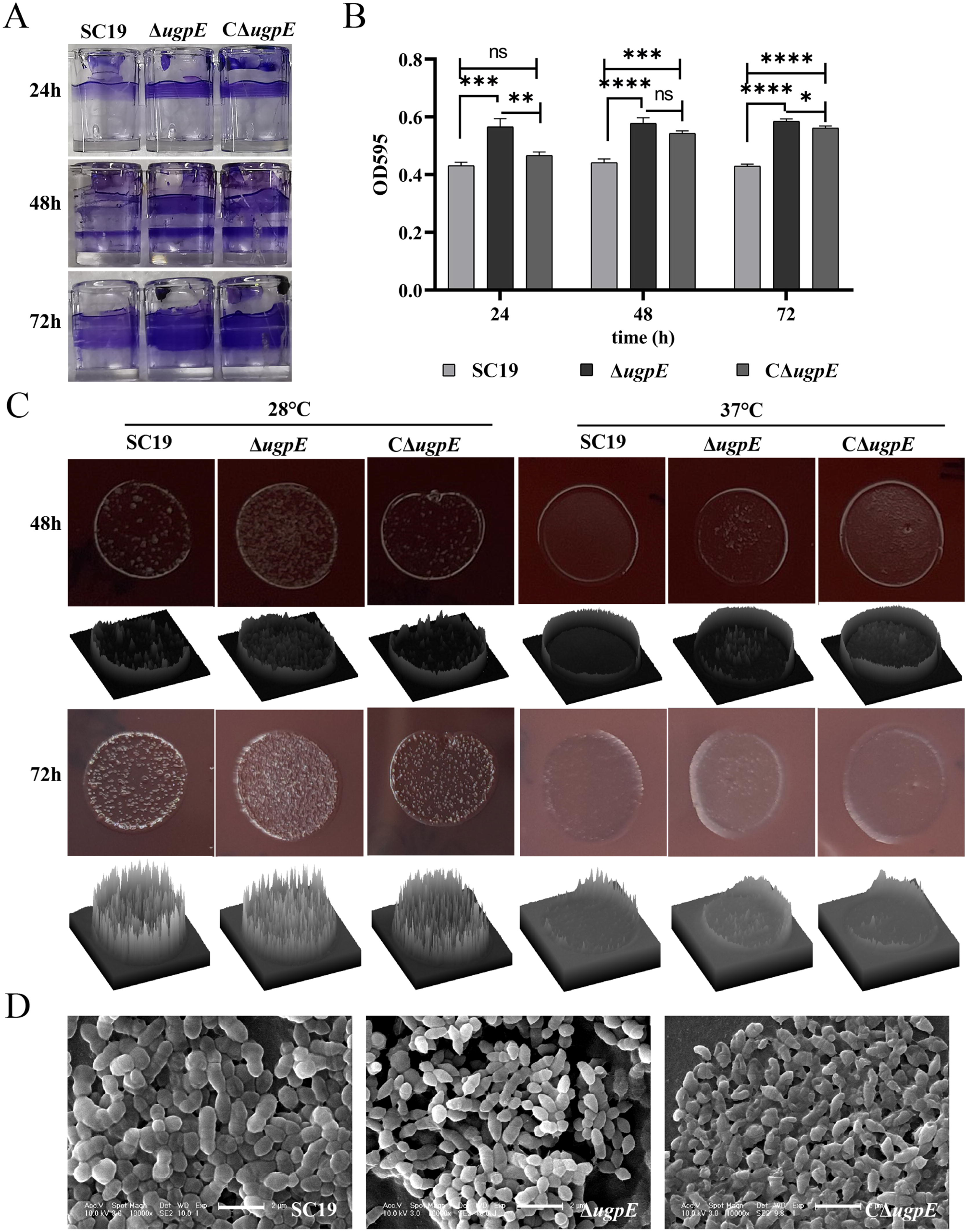
*ugpE* deletion up-regulated SS2 biofilm formation. (A) Observation of biofilms at the air-liquid interface formation on the 96-well plate polystyrene surface after 24h, 48h and 72h of incubation at 37°C. Adherent cells were stained with 0.2% crystal violet. (B) Quantification of biofilms by dissolution in 30% acetic acid and measurement of absorbance at OD595. (C) Analysis of rdar macrocolony morphotype by Congo red agar plate assay after 48h and 72h of incubation at 28°C and 37°C. The surface morphology of the colonies was plotted and analyzed by ImageJ software (1.8.0). (D) Cell morphology analysis by scanning electron microscopy (SEM) of from plate-grown colonies. Size bar, 2 μm. The data are presented as mean ± SD. Data were analyzed using one-way ANOVA (Dunnett’s test). \*\*\**P* < 0.001; \*\**P* < 0.01; \**P* < 0.05. ns, not significant.

Congo red binds to polysaccharides containing contiguous β-(1→4)-linked _D_-glucopyranosyl units and β-(1→3)-_D_-glucans and has been used to identify exopolysaccharides in biofilms [42]. We found that the SC19 wild-type showed a dry and rough morphotype on Congo red agar plates after 48 h and 72 h at 28°C, whereas it was a light red and smooth colony at 37°C (Fig. 3C), suggesting that SC19 produces more extracellular matrix at 28°C than at 37°C. Notably, a drier and rougher morphotype was observed in the Δ*ugpE* mutant compared to the SC19 wild-type and the revertant strain at both temperatures (Fig. 3C), suggesting *ugpE* plays a role in the production of the extracellular matrix of SS2. To our knowledge, this is the first description of regulation of the SS2 rdar morphotype at the gene level. In agreement with the colony morphotype, SEM observation of the agar-grown colonies demonstrated that the production of extracellular matrix was upregulated in the Δ*ugpE* mutant compared to the SC19 wild-type and revertant strains (Fig. 3D).

### ugpE regulates virulence in vitro

As *ugpE* contributes to the capsule production of SS2 (Fig. 1F-H), we assumed that the virulence phenotypes of SS2 might also be affected. We first checked hemolysin activity and found that the hemolysin ability of the Δ*ugpE* mutant was significantly decreased compared to that of the SC19 wild-type and revertant strains (Fig. 4A).

**Fig. 4.**
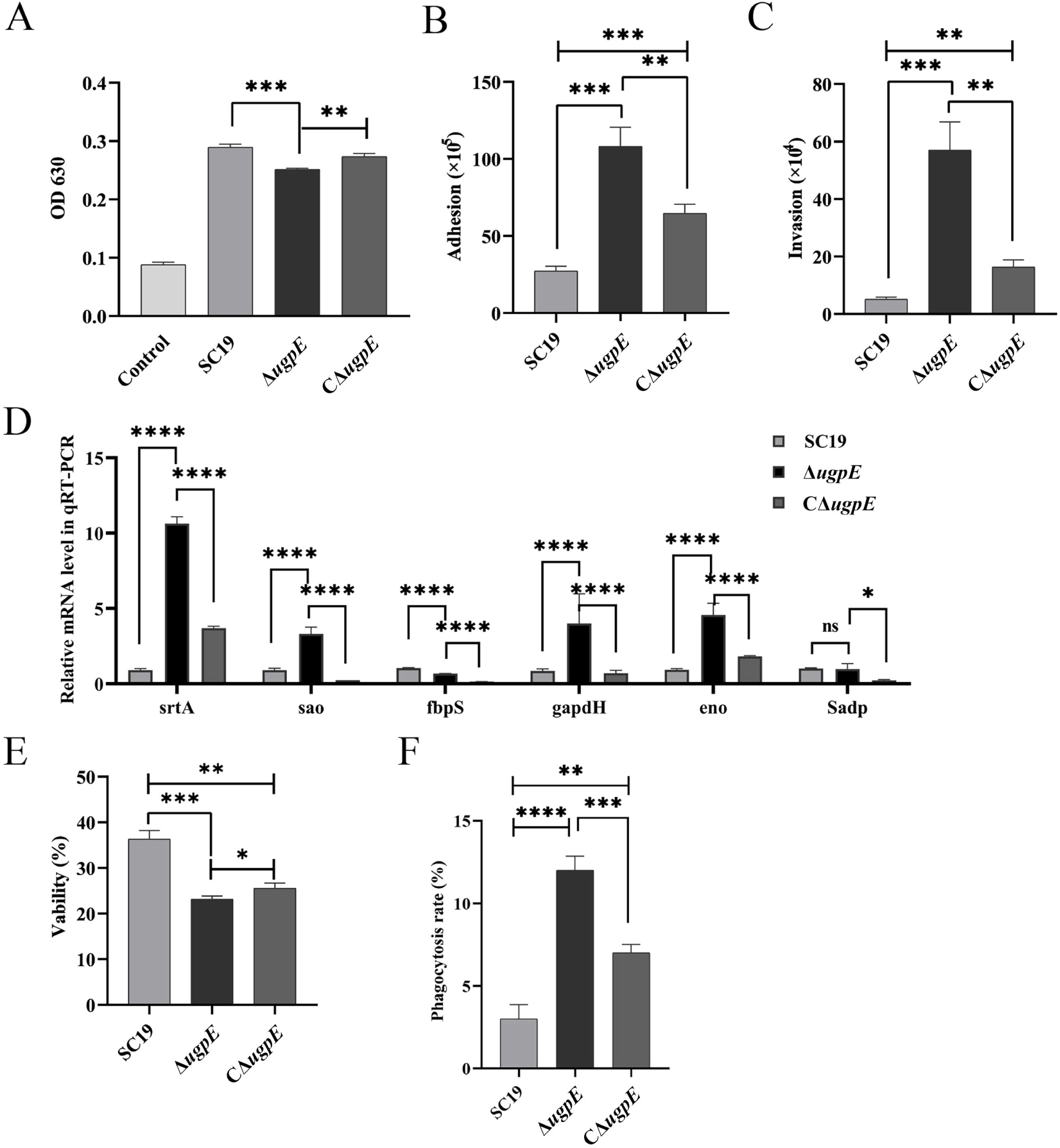
*In vitro* virulence after *ugpE* deletion. (A) Analysis of the hemolysin activity of SC19 wildtype, Δ*ugpE* and CΔ*ugpE* in 5% sheep blood. (B-C) Analysis of adhesion (B) and invasion (C) abilities of each group of cells to hCMEC/D3. (D) Analysis of mRNA levels of adhesion related genes *srtA*, *sao*, *fbpS*, *gapdh*, *eno*, and *Sadp* by qRT-PCR. (E) Survival in whole human blood of each group of cells. Cells were incubated with germ-free human whole blood at 37°C for 2h. (F) Assessment of anti-phagocytosis ability of each group of cells in porcine alveolar macrophages (PAM). The data are presented as mean ± SD. Data were analyzed using one-way ANOVA (Dunnett’s test). \*\*\**P* < 0.001; \*\**P* < 0.01; \**P* < 0.05. ns, not significant.

The ability of bacteria to adhere to and invade host cells is particularly important for bacterial pathogenesis. Subsequently, we investigated the adherence and invasion of each group of strains into human microvascular endothelial cells (hCMEC/D3). Compared to the SC19 wild-type and revertant strains, we found that the number of both adhered and invaded Δ*ugpE* mutant cells to hCMEC/D3 was significantly higher (Fig. 4B and C). With this finding, the transcription levels of adhesion-related genes, including *srtA*, *sao*, *gapdh,* and *eno* were also increased compared to the SC19 wild-type and revertant strains, while the levels of *fbps* and *Sadp* were decreased and remained unchanged, respectively (Fig. 4D). These results indicate that *ugpE* represses adhesion and invasion of host cells.

Bacterial survival in host blood and anti-phagocytosis ability are critical for immune evasion and the establishment of infection. We further assessed these abilities using whole-blood survival and PAM-mediated phagocytosis analyses. Compared to the SC19 wildtype, of which approximately 36% have survived after whole blood killing, the viability of the Δ*ugpE* mutant and the revertant strain declined to 21% and 25%, respectively (Fig. 4E). In line with this result, we found that the phagocytosis rate of the Δ*ugpE* mutant by PAM cells was significantly higher than that of the SC19 wild-type and revertant strains (Fig. 4F), suggesting that the decreased survival ability of the Δ*ugpE* mutant in human blood was possibly due to an increased susceptibility to PAM phagocytosis.

### ugpE is required for SS2 full virulence in vivo

To determine the effect of *ugpE* on the virulence of SS2, an SS2 infection model was constructed in BALB/c mice, and the survival of the mice was assessed. The results showed that all the mice died within 3 days of infection by the SC19 wildtype, while no mice or only half of the mice died of infection by the Δ*ugpE* mutant and the revertant strain CΔ*ugpE*, respectively (Fig. 5A), suggesting *ugpE* is required for SS2 virulence. Furthermore, mice were infected with lower doses of the strains, and the clinical symptoms, weight changes, pathological changes, and bacterial load in the blood and organs were collected and analyzed. Compared to the SC19 wild-type and CΔ*ugpE* infection groups, mice infected with the Δ*ugpE* mutant had significantly lower clinical scores after 12h of infection (Fig. 5B), indicating dramatically relieved virulence of SS2 upon *ugpE* deletion. In terms of body weight, the three groups of infected mice showed similar trends, but the weight loss in the Δ*ugpE* mutant infection group was significantly lower than that in the SC19 wild-type and revertant strains (Fig. 5C). Moreover, the bacterial load in the blood and organs including the brain, liver, kidney, and lung, except in the spleen, was significantly reduced after 24h of infection with the Δ*ugpE* mutant compared to the SC19 wild-type (Fig. 5D), indicating that the colonization ability of SC19 in mice was reduced after *ugpE* deletion, and it was easier to be cleared by the host immune system.

**Fig. 5.**
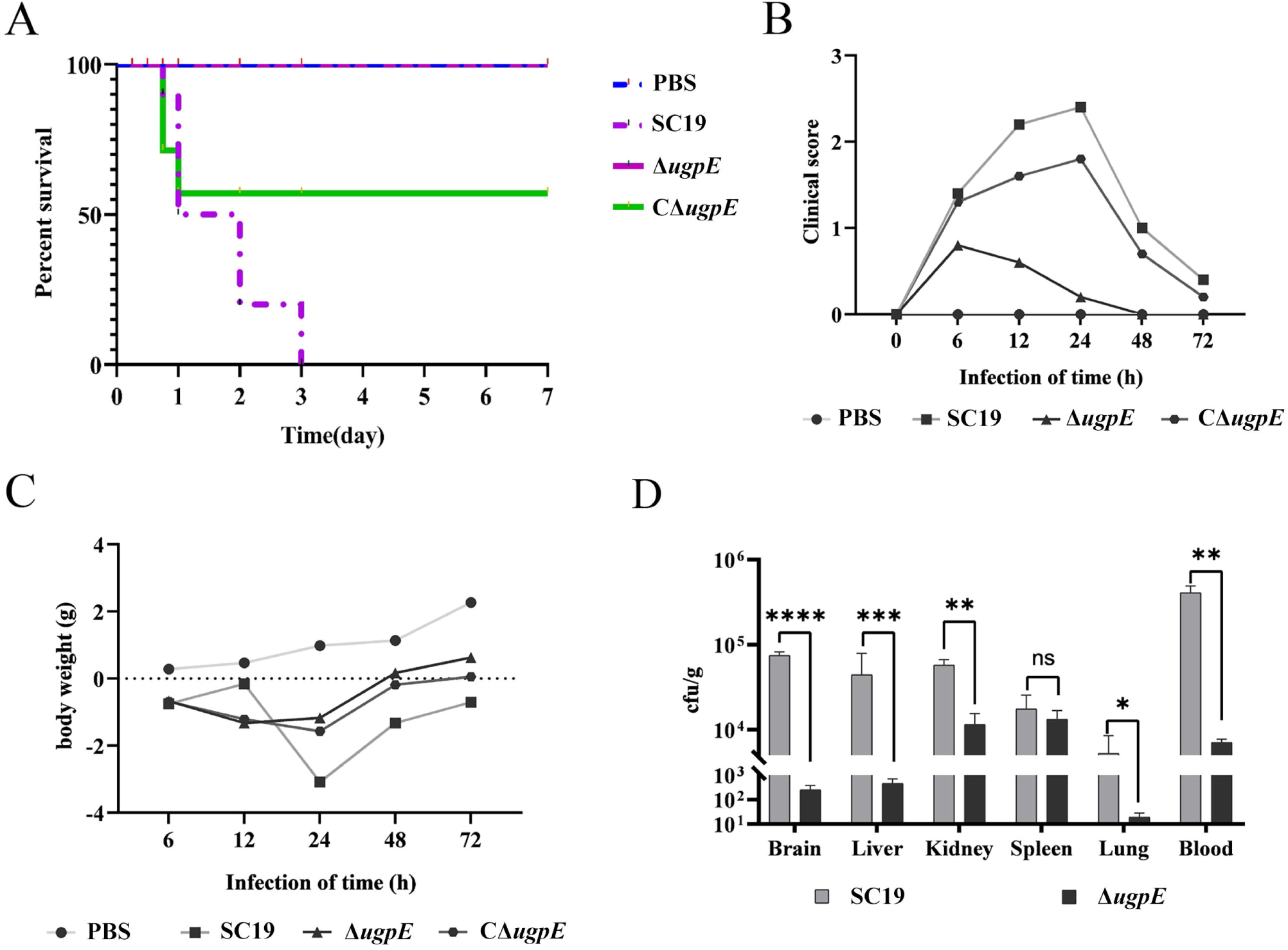
Determination of bacterial virulence after *ugpE* deletion in mice. (A-C) Survival curves (A), clinical scores (B), and body weight changes (C) of the mice infected with SC19 wildtype, Δ*ugpE* and CΔ*ugpE*. (D) Bacterial counts in the major organs and blood of mice after infection with each group of cells. The data are presented as mean ± SD. Data were analyzed using paired *t* tests. \*\*\**P* < 0.001; \*\**P* < 0.01; \**P* < 0.05. ns, not significant.

In line with these observations, the histopathological analysis of each organ revealed that the Δ*ugpE* mutant infected mice showed weaker pathological changes compared to mice infected with wild-type SC19 and CΔ*ugpE* (Fig. 6). Specifically, infection with SC19 wild-type and CΔ*ugpE* led to infiltration of inflammatory cells, including monocytes and neutrophils, in all tissues, especially in the lungs, kidneys, brain, and spleen. Moreover, edema and hyperemia were observed in these tissues upon SC19 wild-type and CΔ*ugpE* infection. In contrast, infection with the Δ*ugpE* mutant alleviated the degree of infiltration of inflammatory cells, edema, and hyperemia, especially in the lungs and brain (Fig. 6), indicating that deletion of *ugpE* impaired the virulence of SC19. Taken together, these results demonstrated *ugpE* is required for the full virulence of SS2 in mice.

**Fig. 6.**
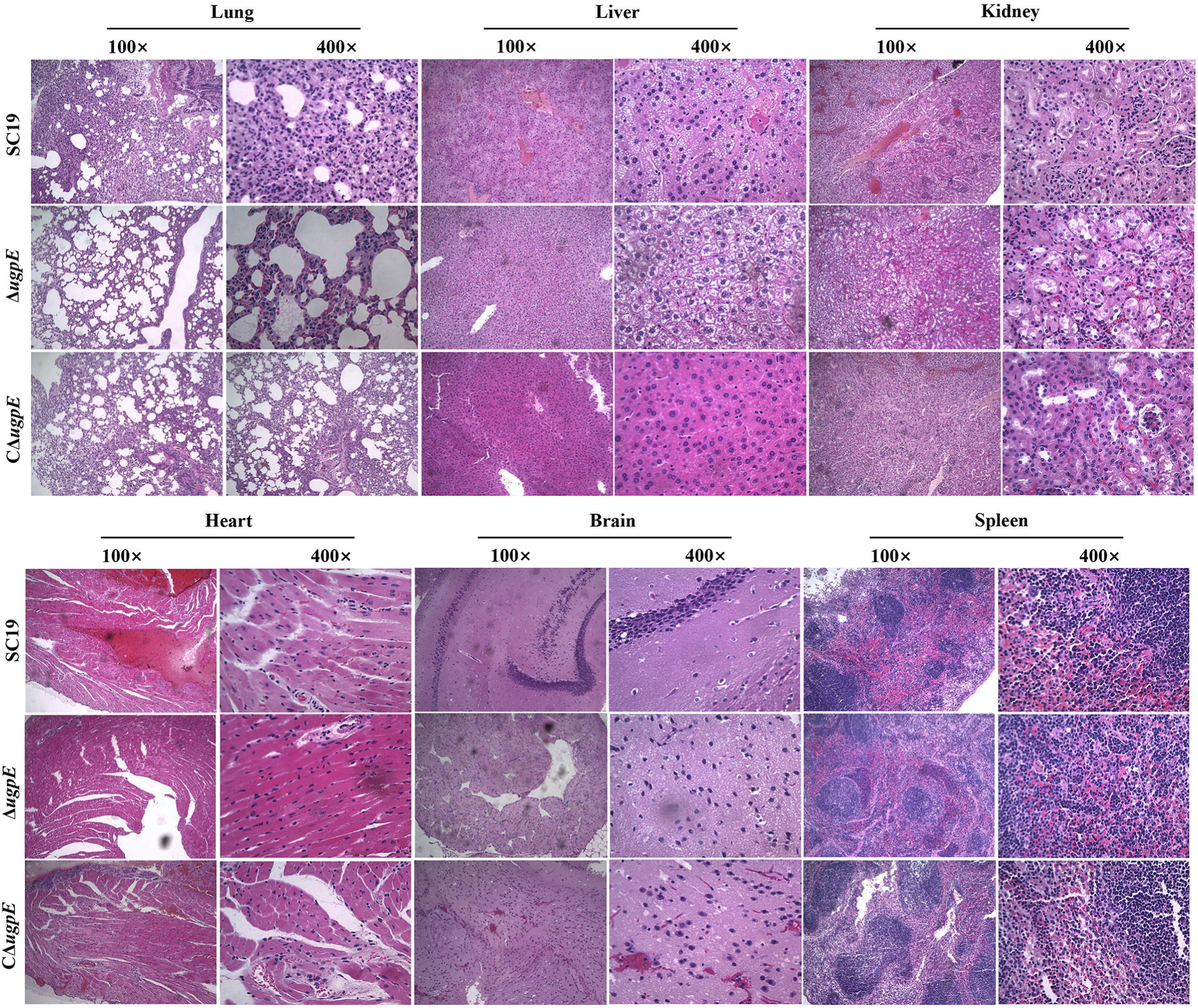
Pathological changes in lung, liver, kidney, heart, brain, and spleen upon infection with SC19 wildtype, Δ*ugpE* and CΔ*ugpE* strains using histopathological analysis. The tissue slices were stained with hematoxylin and eosin (HE) and the histopathological changes were indicated with black arrows.

### ugpE deletion alters distribution of macrophages in major organs of mice upon infection

Since *ugpE* is required for full virulence both *in vitro* and *in vivo*, we wondered whether the host immune response was also altered upon SS2 infection. To this end, the distribution of macrophages in the major organs of mice upon SS2 infection was assessed using flow cytometry. The results showed that the percentage of macrophages in the lung after 24h infection with the Δ*ugpE* mutant was significantly lower than that after infection with the SC19 wildtype and CΔ*ugpE* (Fig. 7A and B). However, the percentage of macrophages in other organs after 24h infection of with the Δ*ugpE* mutant was not significantly different from the other two groups (Fig. 7A, C-F).

**Fig. 7.**
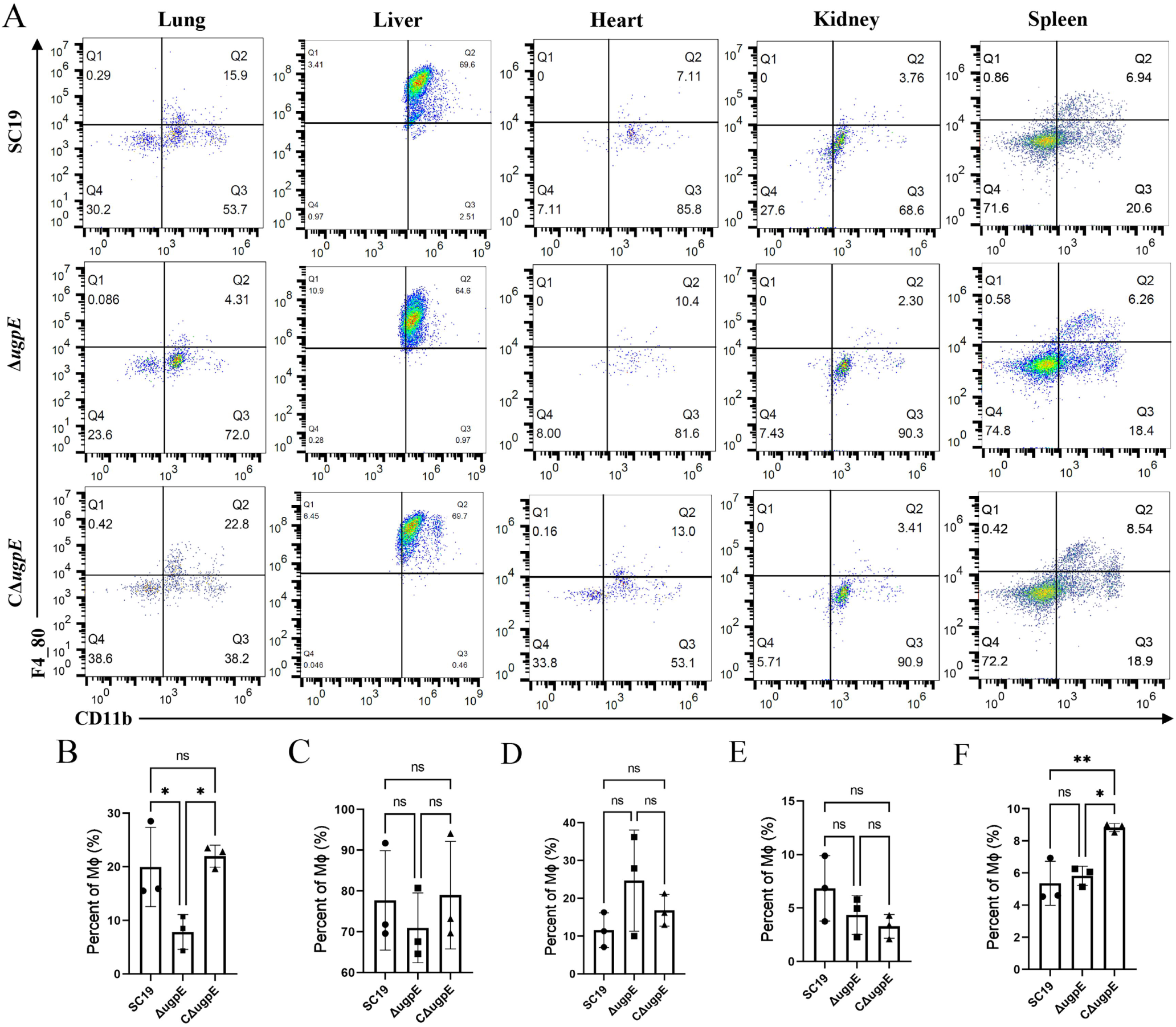
Distribution of macrophages in major organs of mice upon infection with SC19 wildtype, Δ*ugpE* and CΔ*ugpE* strains by flow cytometry. The data are presented as mean ± SD. Data were analyzed using one-way ANOVA (Dunnett’s test). \*\*\**P* < 0.001; \*\**P* < 0.01; \**P* < 0.05. ns, not significant.

## Discussion

UgpABCE is a member of the ABC transporter superfamily that is involved in glycerophospholipid synthesis in *E. coli* [23]. In this study, we report a novel role of the Ugp transporter subunit UgpE in the regulation of various cellular phenotypes, including stress tolerance, biofilm formation, and virulence of the zoonotic pathogen *S. suis*. Deletion of *ugpE* impaired cell chain formation, capsular synthesis, and stress tolerance while upregulating biofilm formation and adhesion to hCMEC/D3 cells. Importantly, bacterial virulence was attenuated in mice after deletion of *ugpE*. These results suggest *ugpE* is a novel virulence factor that regulates the various cellular phenotypes of *S. suis*.

UgpE is a subunit of the UgpABCE transporter system. This system is predicted to be prevalent in diverse bacteria [43] but has been well studied in *E. coli* and several other bacterial species only, including *Mycobacterium tuberculosis* [44] and *Thermus thermophilus* [45]. To the best of our knowledge, this is the first study to identify an UgpE homolog in *S. suis*. In *E. coli*, UgpE contributes to the transport of both G3P and glycerophosphodiesters from the periplasm to cytosol under phosphate starvation conditions [23, 44]. Glycerophosphodiesters are degraded into G3P and further acylated by *sn*-glycerol-3-phosphate acyltransferase to initiate the biosynthesis of membrane glycerophospholipids or central carbon metabolism [46, 47]. Indeed, glycerophosphodiesters are also taken up by Glp (<u>gl</u>ycerol-3-<u>p</u>hosphate transporter), apart from the UgpABCE transporter system [48]. However, preliminary analysis demonstrated that no Glp homolo was present in the SC19 genome (data not shown), indicating that the UgpABCE transporter might be the sole system for G3P and glycerophosphodiester acquisition in SC19. Interestingly, we did not observe any obvious morphological changes in the cell membranes or cell walls among the groups (Fig. 1F and H). Considering the importance of glycerol phosphate in membrane phospholipid synthesis, we hypothesized that other cell wall components might be impaired by *ugpE* deletion.

In addition to participating in glycerophospholipid biosynthesis, glycerophosphodiesters also serve as osmoprotectants in eukaryotic cells. In *Saccharomyces cerevisiae*, glycerophosphocholine produced from the membrane lipid phosphatidylcholine protects cells from hypersaline stress [25]. In line with this phenomenon, the Δ*ugpE* mutant dramatically diminished the anti-stress abilities of SC19 compared to the wild-type SC19 (Fig. 2), suggesting *ugpE* is required for bacterial stress responses to adverse environmental conditions. Although the cell membrane and cell wall were hardly affected, the capsule structure almost disappeared (Fig. 1F-G). The capsule is a protective polysaccharide layer around the cell, which is an important virulence factor for most encapsulated streptococci. The absence of CPS correlates with increased phagocytosis by macrophages and decreased virulence in murine and pig models of infection [49]. In SS2, the capsular locus (*cps*) responsible for CPS biosynthesis involves 25 genes [40]. It has been shown that inactivation of the *cps2B* gene for polysaccharide chain length determination, *cps2EF*, *cps2G*, *cps2J* and *cps2L* genes for glycosyltransferases leads to impaired capsule production [7, 50]. *cpsBCD* genes form a tyrosine phosphoregulatory system that controls the CPS assembly machinery, of which CpsB assists in CpsD dephosphorylation and its interaction with CpsC, leading to elevated CPS polymerization [51, 52]. The *cpsPQRST* genes showed homology to the NeuBCDA proteins of Group B *Streptococcus* (GBS) and *E. coli* and thus have been predicted to be involved in the synthesis of sialic acid (NeuNAc), one of the main components of the CPS of *S. suis* [40]. Inactivation of *neuC* (*cpsQR*) resulted in the loss of the whole capsule of *S. suis*, decreased the anti-phagocytosis ability of macrophages, and impaired virulence in mice [53]. In GBS, *neuD* functions as a Sia O-acetyltransferase and is required for the synthesis of the unique sialylated CPS shared only by *S. suis* and GBS [54, 55]. In this study, a reduction in the transcription levels of *cps-2B*, *cps-2C* and *cps-2S* was observed in the Δ*ugpE* mutant (Fig. 1I), suggesting that the unencapsulated phenotype of the Δ*ugpE* mutant may be due to interference with polysaccharide chain formation and the expression of sialic acid for CPS synthesis in *S. suis*. In line with these observations, the unencapsulated Δ*ugpE* mutant exhibited a lower survival rate in whole human blood, antiphagocytic ability against PAM, and decreased virulence in mice compared to the SC19 wild-type (Fig. 4E-F; Fig. 5), suggesting that deficiency in CPS development is a critical reason for the weakened virulence of *S. suis* by *ugpE* deletion.

Moreover, only three primary capsule synthesis pathways have been identified so far. The Wzx/Wzy-dependent and synthase-dependent pathways are present in both gram-positive and gram-negative bacteria, while the ABC transporter-dependent pathway is present only in gram-negative bacteria [56, 57]. Capsule synthesis is regulated by small RNAs (sRNA). In *S. suis*, sRNA rss04 contributes to meningitis induction in mice by regulating CPS synthesis [58]. In this study, we identified for the first time that UgpE, which belongs to the Ugp transporter system, regulates CPS synthesis in Gram positive streptococci.

Notably, both the adhesion and invasion abilities of host hCMEC/D3 cells were enhanced in the Δ*ugpE* mutant compared to those in the SC19 wild-type (Fig. 4B-C). We speculated that although CPS is a critical antiphagocytic factor that resists phagocytosis by macrophages and neutrophils in the blood, it may interfere with the adhesion and invasion of host cells by hindering cell wall components, especially some vital virulence factors located on the cell surface for adhesion. Indeed, many studies have demonstrated that the repression of CPS production enhances the adhesion and invasion of SS2 to host cells and impairs virulence in mice [58, 59, 60]. As anticipated, the transcription levels of genes encoding cell surface virulence factors, including *srtA*, *sao*, *gapdh,* and *eno* were upregulated in the Δ*ugpE* mutant compared to those in the parental strain (Fig. 4D). These results suggest that impairment of CPS by the Δ*ugpE* mutant may lead to exposure of cell surface proteins, which contributes to bacterial adhesion and invasion of host cells. In contrast, impairment in CPS decreased antiphagocytic ability against macrophages and neutrophils in the blood, resulting in attenuated virulence, as confirmed by the mouse infection model (Fig. 5; Fig. 6).

Biofilm formation represents a protected mode of growth that renders bacterial cells less susceptible to host immune responses and thus enables pathogens to survive in hostile environments [41]. Interestingly, the Δ*ugpE* mutant exhibited enhanced biofilm formation on the polystyrene surface, along with decreased survival in whole human blood and anti-phagocytic ability in PAM (Fig. 3A-B; Fig. E-F). Considering the critical role of CPS in the resistance to phagocytosis, we reasoned that infection by the Δ*ugpE* mutant may not be established due to the loss of CPS structure, even through the upregulation of biofilm formation. Indeed, studies have shown that the CPS structure affects bacterial biofilm formation. In *S. pneumoniae*, encapsulated clinical pneumococcal isolates are impaired in their capacity to form biofilms, and reduction in the amount of CPS is usually correlated with increased biofilm-forming capacity [61]. In SS2, the flavonoid compound rutin specifically downregulated biofilm formation by interfering with CPS biosynthesis [62].

In summary, our study reported for the first time that UgpE, which belongs to the Ugp transporter system, is required for full virulence and biofilm formation of the zoonotic pathogen *S. suis*. UgpE contributes to bacterial virulence by positively regulating CPS synthesis, demonstrating that UgpE is a novel virulence factor of *S. suis*. Our results expand the current knowledge on the pathogenesis of *S. suis*.

## Ethical statement

Animal infection experiments were conducted at the Laboratory Animal Center of College of Veterinary Medicine, Jilin University (Permit number: KT202103086) in accordance with the Regulations for the Administration of affairs Concerning Experimental Animals.

## Authors’ contributions

**Qiulei Yang**: Methodology, Data curation, formal analysis, Investigation, Writing–original draft. **Na Li**: Methodology, Data curation, formal analysis, and investigation. **Yanyan Tian**: Methodology, Investigation. **Qiao Liang, Miaomiao Zhao, Hong Chu**: Validation, Visualization. **Yan Gong, Tong Wu, Shaopeng Wei, He Wang**: Investigation. **Guangmou Yan**: Resources. **Fengyang Li**: Conceptualization, Funding acquisition, Supervision, Writing–original draft, writing–review, and editing. **Liancheng Lei**: Funding acquisition, Supervision, Project administration, Resources, Writing, review, and editing.

## Data availability statement

All data generated or analyzed during this study are deposited in Figshare (10.6084/m9.figshare.27094336).

## Acknowledgements

We thank Prof. Anding Zhang (Huazhong Agricultural University, Wuhan, China) for providing *S. suis* serotype 2 (SS2) strain SC19, and Prof. Yan Chen (Jilin University, Changchun, China) for providing hCMEC/D3 cell line.

